# Matrix stiffness induces endothelial network senescence

**DOI:** 10.1101/2025.10.05.680536

**Authors:** Jiyeon Song, Alexandra N. Rindone, Ya Guan, Prarthana Sanjay Daswani, Jennifer H. Elisseeff, Sharon Gerecht

## Abstract

Identifying the drivers of cellular senescence that contribute to the decline in tissue function related to aging- and disease is critical for developing restorative interventions. Here, we investigated how increased mechanical stress from extracellular matrix (ECM) stiffening shapes endothelial cell (EC) senescence. We developed a 3D human *in vitro* model that decouples mechanical stress from inflammatory or biochemical inputs, enabling the study of senescence responses to tissue stiffening alone. We found that matrix stiffening induces an EC senescence phenotype with elevated p16/p21 and an immunomodulatory senescence-associated secretory phenotype (SASP), in the absence of inflammatory signals. This mechano-induced senescence state engaged a Notch-JNK-FOS signaling axis, and pharmacologic inhibition of Notch attenuated stiffness-induced senescence. Supporting the translational relevance of this mechanism, analysis of fibrotic capsule tissue from patients with synthetic breast implants, a model of localized, mechanically driven fibrosis, revealed increased p16^+^Notch1^+^ endothelial populations. Complementary single-cell RNA sequencing data confirmed their enrichment in Notch/JNK- and SASP-related gene programs. Together, these findings define vascular senescence as a mechanosensitive process and identify tissue stiffening as an upstream aging signal. Our work offers a human-relevant platform for studying targetable stages of endothelial mechanoaging.

## Main

The extracellular matrix (ECM) is a dynamic regulator of cell fate, transmitting mechanical signals that shape cell behavior. Progressive matrix stiffening occurs in aging and diseases, due to increased matrix deposition and crosslinking, ultimately altering the mechanical forces sensed by surrounding cells^1^. This shift disrupts homeostatic cell-matrix communication, driving changes in signal transduction, gene expression, and cellular phenotypes.^2,3^ These effects are especially evident in the microvasculature, where stiffened and disorganized ECM accumulates around endothelial cells (ECs), promoting a shift toward a more contractile and dysfunctional state.^4^ This EC transition compromises adherens junctions and weakens barrier integrity.^5^ However, despite clear evidence of these tissue-level changes, the cellular-level consequences of altered mechanical cues remain poorly understood, particularly whether matrix stiffening triggers phenotypic transitions to senescence.

While many studies have focused on biochemical inducers of EC senescence, such as oxidative stress, inflammation, and chemotherapeutic agents,^6^ the role of mechanical stress remains unclear. ECM stiffening is an inevitable feature of aged or fibrotic tissue, and biomechanical stress is continuously imposed on the microvasculature. However, whether such biomechanical forces alone induce EC senescence remains an open question, due in part to the difficulty of isolating mechanical cues in complex *in vivo* tissue environments^7,8^.

Here, we developed a tunable hydrogel platform that models the dynamic increase in ECM stiffness to investigate how mechanical forces regulate EC network senescence and its underlying molecular mechanisms. This engineered system captured a mechanically induced, transitional senescence state in ECs, marked by elevated senescence marker expression and an immunomodulatory senescence associated secretory phenotype (SASP), distinct from the classical IL-6/IL-8-enriched profile. Mechanistically, this mechanoaging response was driven by a Notch-JNK-FOS signaling axis that promoted a FOS-dominant AP-1 program, independent of canonical c-JUN nuclear activation. This highlights a mechanosensitive senescence pathway that remains incompletely characterized in the context of vascular aging. Validation of the model with fibrotic capsule tissue from human breast implants and accompanying single-cell RNA sequencing data revealed an enrichment of Notch1^+^ senescent ECs in stiffened environments, supporting the translational relevance of our *in vitro* findings. This work establishes a mechanosensitive platform to dissect biomechanical inputs and identify targetable drivers of endothelial senescence.

## Results

This study aimed to uncover translationally relevant mechanisms by which matrix stiffening drives blood EC network senescence. To this end, we established a physiologically relevant 3D model that supports the formation of luminalized endothelial networks in a soft collagen-rich matrix, followed by on-demand stiffening under controlled conditions. This two-step crosslinking strategy recapitulates the increase in ECM stiffness, while holding biochemical cues constant, thereby isolating mechanics as the independent variable.^4^ Unlike *in vivo*, where stiffness changes coincide with shifts in pathological soluble factors, our system enables selective interrogation of mechanical effects on EC network senescence.

We previously established a methacrylated collagen (Col-MA) and hyaluronic acid (HA-MA) platform in which collagen first forms a physically crosslinked gel at 37°C, followed by ruthenium-mediated, light-induced polymerization of methacrylate groups.^4^ In this first-generation system,^4^ increasing visible light exposure to tune gel stiffness also elevated the flux of photogenerated radicals and reactive oxygen species (ROS) byproduct, confounding mechanical effects with ROS-driven senescence^2^. To prevent this uncontrolled sequel, we developed a second-generation Col-MA/HA-MA system that exhibits different stiffness levels under the same photochemical conditions. Collagen I was retained as the bioactive scaffold for structural and mechanotransducive cues.^9,10^ Hydrogel stiffness was modulated by varying the degree of HA methacrylation (14 and 64 mol%; **Fig. S1**) while holding light intensity, duration, and photoinitiator constant. Under these matched conditions, 14 mol% HA-MA yielded an intermediate (IM) stiffness, whereas 64 mol% produced a stiffer matrix, while the Soft low and high formulations prepared with different HA-MA variants had comparable moduli (**Fig. 1A; Fig. S2A**).

**Figure 1:**
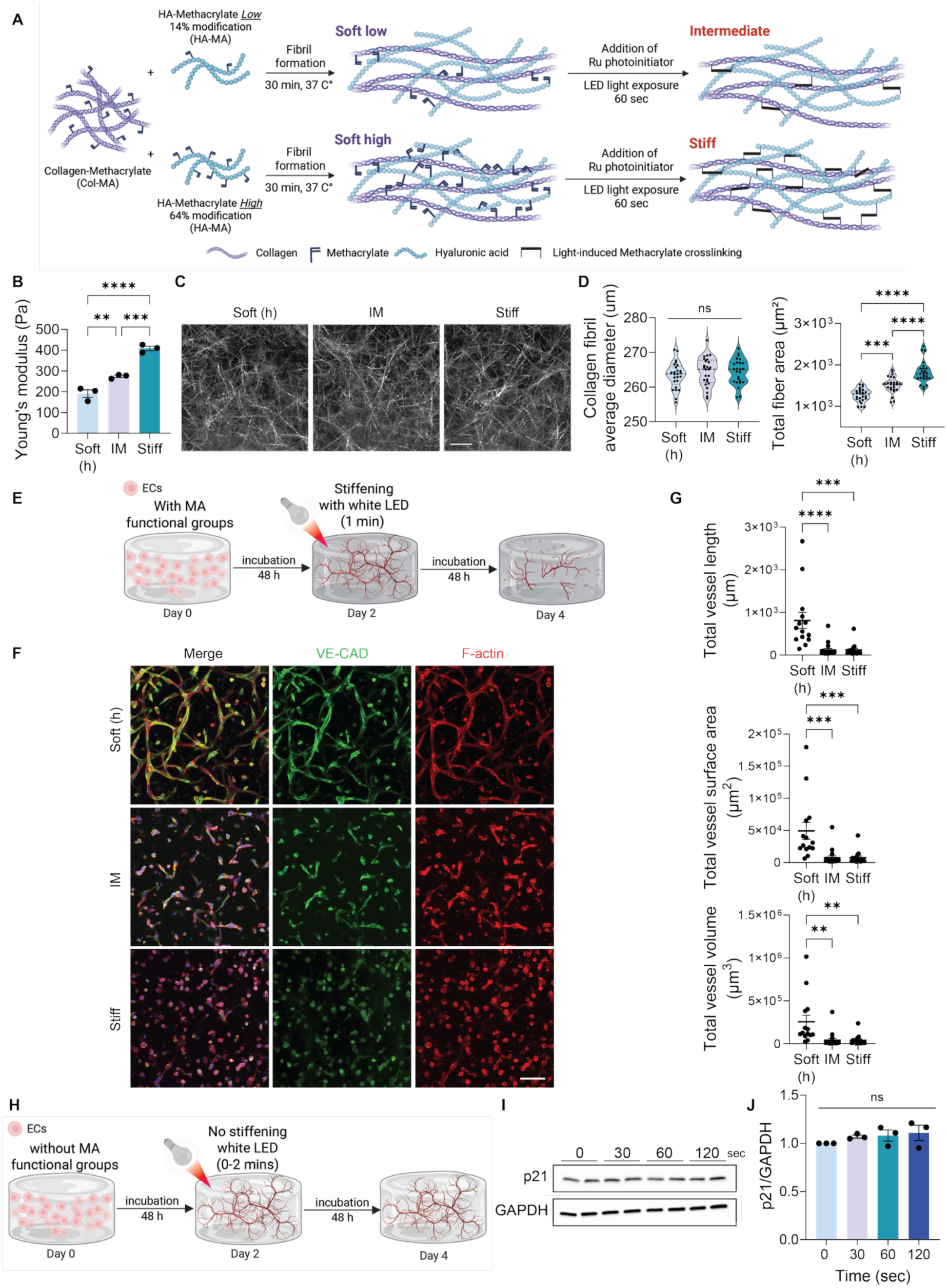
*In vitro* hydrogel system to model endothelial network aging. (A) Schematic of the Col-MA/HA-MA hydrogel system using two different degrees of methacrylate incorporation on HA. (B) Young’s modulus of Soft (high), Intermediate (IM), and Stiff hydrogels. N=3. (C) Representative reflective confocal images of collagen fiber structures at varying stiffness levels. Scale bar= 20 µm. (D) Quantification of collagen fibril average diameter and total fiber area. N=5 with 5 fields of images in each. (E) Schematic of the 3D stiffening hydrogel system. ECs were encapsulated in Col-HA hydrogels and allowed to form networks for 48 h, followed by photo-crosslinking using a ruthenium photoinitiator for 1 min. Both soft control and stiffened hydrogels were subsequently cultured for an additional 48 h prior to analysis. (F) Representative maximum intensity projections of confocal z-stack showing microvascular networks in Soft, IM, and stiff hydrogels. Increased stiffness reduced vessel formation, as quantified in (G) by total vessel length, surface area, and volume. N=2. DAPI in blue, VE-Cad in green, F-actin in red. The scale bars are 100 µm. (H) Schematic of control experiment assessing the effect of radical exposure in the absence of matrix stiffening. ECs were encapsulated in unmodified collagen hydrogels lacking MA functional groups and incubated for 48 h. Cell-laden hydrogel constructs were then exposed to white LED light (0-120 seconds) in the presence of photo initiator, followed by continued culture to Day 4. (I) Western blot analysis and (J) quantifications of p21 levels in ECs encapsulated in unmodified collagen hydrogels (no stiffening) to evaluate the effect of radical exposure on cellular senescence. p21 levels were normalized to GAPDH. N=3. Significance levels were set at *p ≤ 0.05, **p ≤ 0.01, ***p ≤ 0.001, and ****p ≤ 0.0001.

Oscillatory rheometry measurements confirmed effective stiffness separation, with Young’s modulus ∼2.28 fold higher in Stiff and ∼1.43 fold higher in IM relative to Soft (**Fig. 1B**), closely mimicking the change in ECM stiffening in aging mouse tissue compared to young^4^. Equilibrium swelling ratios were 397.01 ± 47.68, 254.62 ± 46.92, and 184.19 ± 7.62 for Soft, IM, and Stiff, respectively (**Fig. S2B**). Reflective confocal imaging revealed unchanged collagen fibril with across groups (263-264 nm) but increased total collagen fiber area with stiffness (**Fig. 1C-D**), consistent with higher crosslink density rather than changes to the intrinsic collagen/HA network structure. The two Soft formulations also exhibited comparable swelling ratios, fibril widths, and fiber areas, indicating differences in HA methacrylation did not alter hydrogel properties prior to matrix stiffening (**Fig. S2A-E**).

We next examined whether microvascular networks regress in response to matrix stiffening, as observed in aged kidney^4^. We encapsulated endothelial colony-forming cells (ECFCs) in the hydrogels and allowed them to form networks for 48 hours. Matrix stiffening was then induced *in situ* with a photoinitiator and LED illumination, followed by culture for an additional 48 hours (**Fig. 1E**). In both the IM and Stiff conditions, disrupted network morphology with significantly reduced total vessel length, surface area, and volume was observed (**Fig. 1F-G**). By contrast, networks in Soft gels remained intact and continued to mature, highlighting stiffness-dependent effects on vascular phenotype and aligning with our previous observation.^4^ Consistently, both Soft formulations without light exposure exhibited comparable microvessel formation, confirming that different levels of methacrylate functionalization did not affect microvasculature formation capacity (**Fig. S2F-G**). Overall, no differences were detected in the material properties and EC network assembly between the two soft hydrogel formulations (**Table S1**). Consequently, we used the soft gel with higher MA modification in the HA as the control in all our experiments. Additionally, since both IM and Stiff hydrogels are polymerized under the same photoinitiation conditions (**Table S1**), any variations in cellular responses can be attributed to matrix stiffness rather than radical exposure. This dynamic stiffening system effectively isolates matrix stiffness as the only variable, allowing us to study EC senescence due to mechanics rather than ROS generated during photopolymerization.

To further ensure that senescence outcomes reflected mechanics rather than radical exposure, we tested whether photoinitiator-derived radicals alone induce EC senescence in the absence of matrix stiffening. We used unmodified collagen hydrogels, which do not stiffen upon photoinitiator and light exposure, and subjected ECs to light for 0-120 seconds, spanning durations shorter and longer than our standard protocol (**Fig. 1H**). p21 senescence marker expression was unchanged across time points (**Fig. 1I-J**), indicating that, at the doses used here, radical exposure alone does not trigger senescence. Therefore, by varying stiffness under identical photochemical conditions, our *in vitro* stiffening hydrogel model allows for a mechanics-specific readout of EC network senescence.

**Figure S1:**
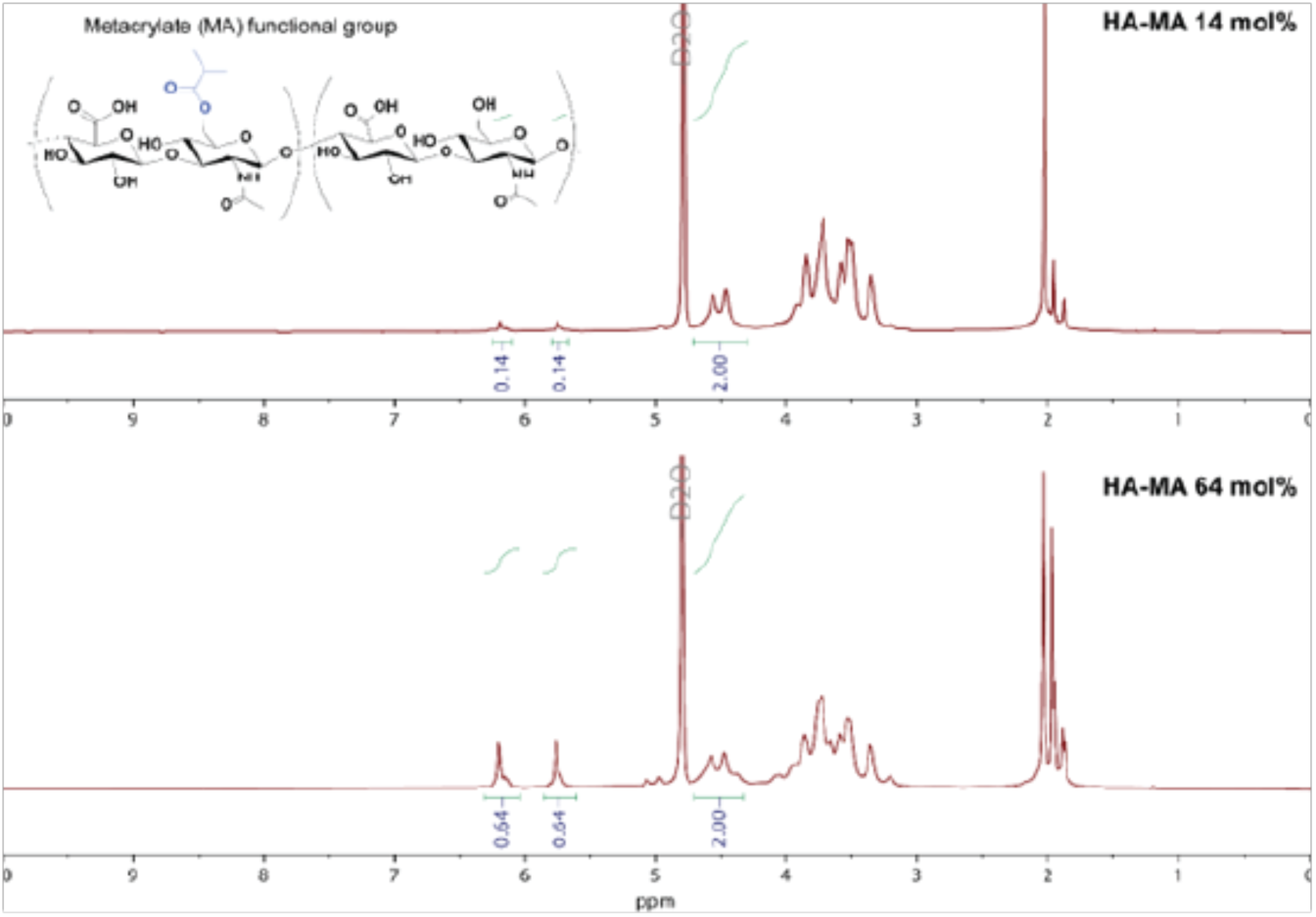
^1^H NMR spectra for HA-MA low (14 mol%, top) and high (64 mol%, bottom) in D_2_O.

**Figure S2:**
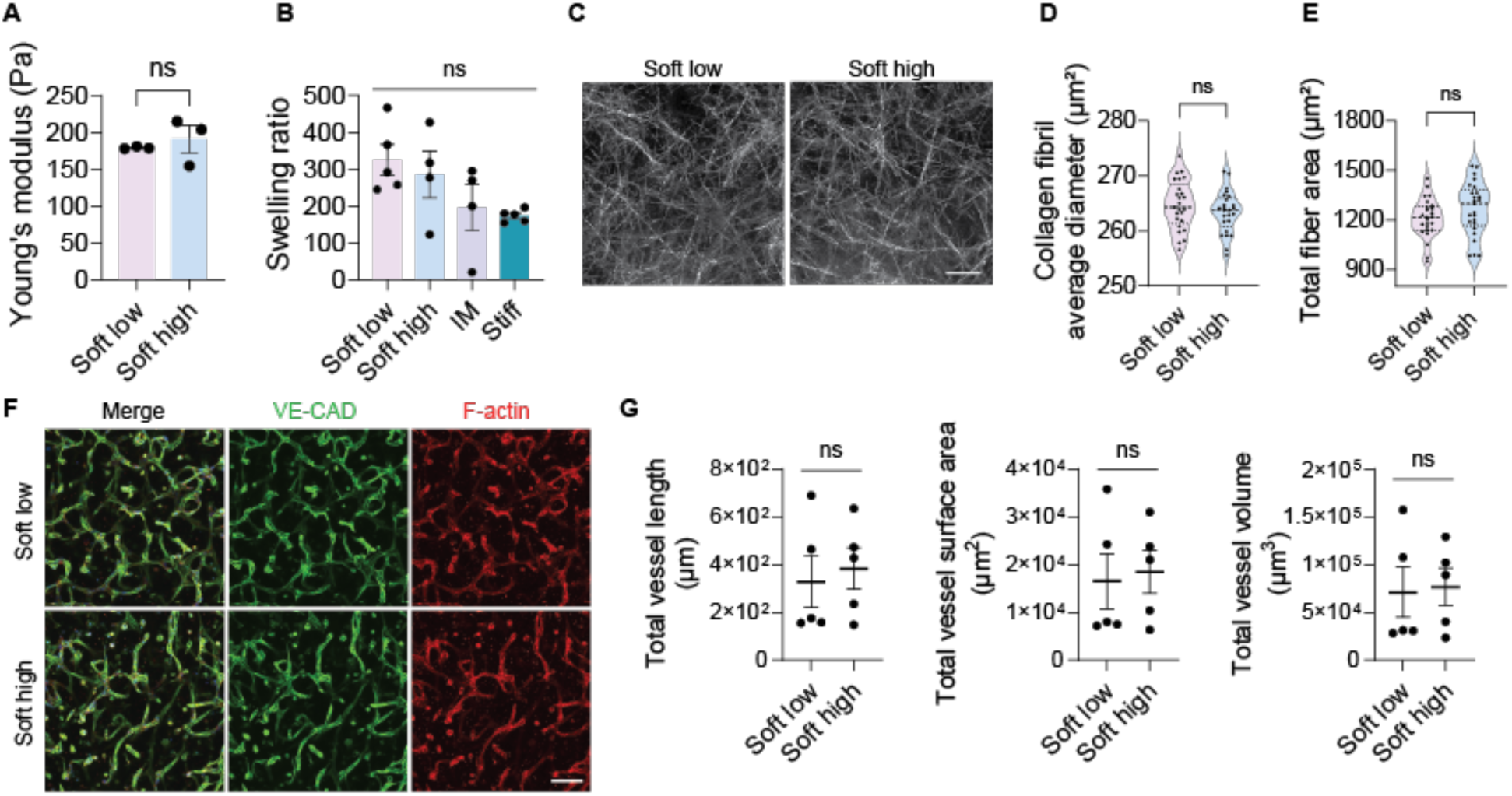
Hydrogel characterization of two Soft formulations prepared with varying methacrylate incorporation on HA. (A) Young’s modulus measured by rheometry. (B) Equilibrium swelling ratio. (C) Representative confocal reflectance images showing comparable fibrillar microstructure (scale bar: 20 μm) as quantified in (D) by collagen fibril average diameter and in (E) by total fiber area. N=5 with 5 fields of images in each. (F) Representative maximum intensity projections of confocal z-stack showing microvascular networks in Soft low and Soft high hydrogels (Scale bar: 100 µm). (G) Quantification showing comparable microvascular network formation in terms of total vessel length, surface area, and volume.

**Table S1.**
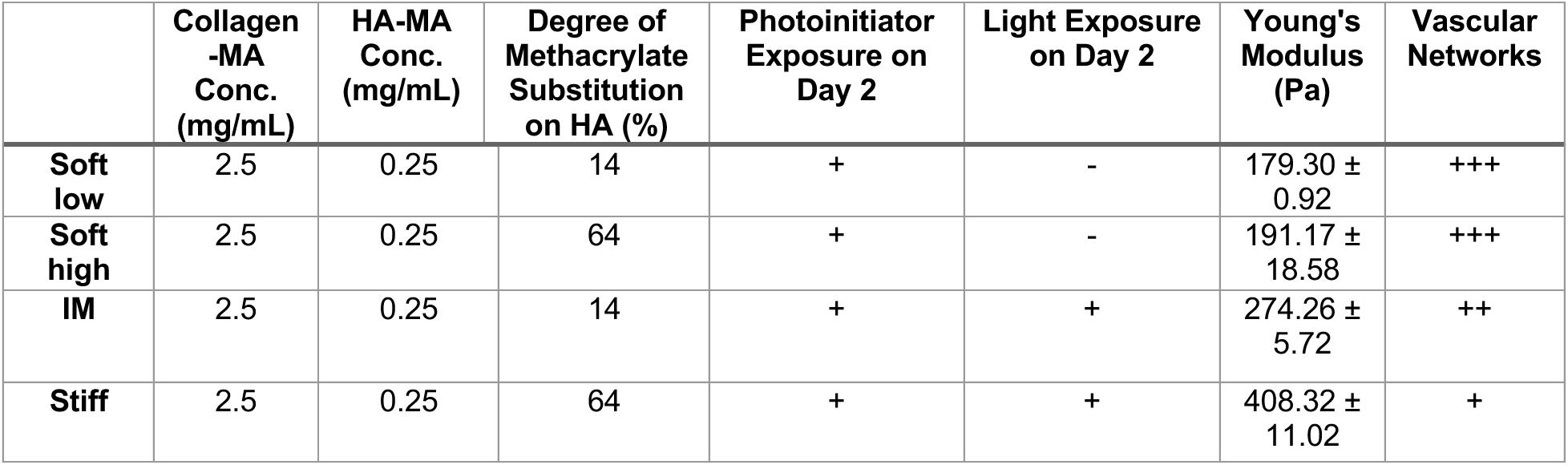
Summary of hydrogel formulations and secondary crosslinking parameters used to model dynamic matrix stiffening. Conditions vary by methacrylation level and light exposure to achieve a tunable range of stiffness, as reflected in Young’s modulus measurements.

Next, we tested whether dynamic matrix stiffening drives endothelial network senescence. On day 2, endothelial networks were exposed to stiffened matrix environments corresponding to IM and Stiff conditions. At the transcription level, the senescence markers *CDKN1A* (p21) and *CDKN2A* (p16) were significantly upregulated with increasing matrix stiffness, with the highest expression level observed in Stiff gels (**Fig. 2A**). Consistently, Western blot showed a stiffness-dependent increase in p21 protein (**Fig. 2B-C)**. Additional hallmarks, p16 and SA-β-Gal activity assessed by immunofluorescence were elevated in Stiff matrices (**Fig. 2D-G**), indicating that matrix stiffening promotes EC senescence.

**Figure 2:**
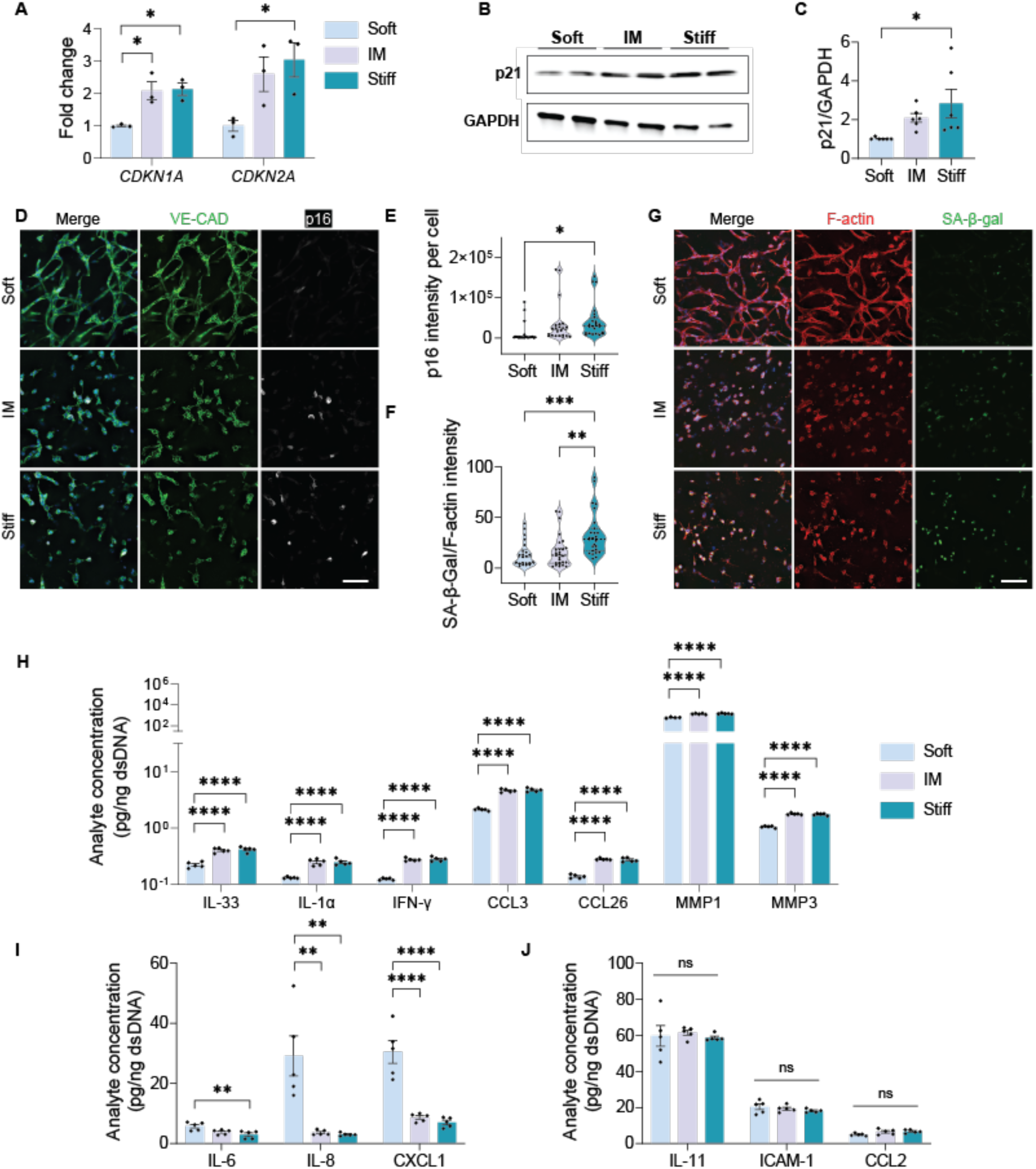
Matrix stiffness-induced endothelial network senescence. Endothelial networks cultured in hydrogels of increasing stiffness show elevated expression of senescence markers: (A) *CDKN1A* and *CDKN2A* mRNA levels measured by qRT-PCR, (B-C) p21 protein levels assessed by Western blot, and (D-E) p16 protein expression visualized by immunofluorescence staining. Representative confocal images show microvascular networks stained for VE-Cadherin in green, p16 in while, nuclei in blue. Scale bar is 100 µm. N=3. (F-G) SA-β-Gal activity increased with matrix stiffening. Representative confocal images show networks stained for F-actin (red), SA-β-Gal (green), and nuclei (blue). Scale bar: 100 µm. N=3. (H-J) Senescence-associated secretory phenotype (SASP), including cytokines, chemokines, and MMPs analyzed using a Luminex assay. Values were normalized to the amount of dsDNA isolated from each hydrogel construct. N=3. Significance levels were set at ns= not significant (p > 0.05), **p ≤ 0.01, and ****p ≤ 0.0001.

In addition to upregulating senescence markers, dynamic matrix stiffening also induced a distinct senescence-associated secretory phenotype (SASP). To characterize this mechanoaging-associated SASP, we analyzed culture supernatants for cytokines and proteases secreted by senescent ECs in the absence of exogeneous inflammatory stimulation. Luminex analysis revealed that matrix stiffening significantly increased secretion of several immunomodulatory molecules, including IL-33, IL-1α, IFN-γ, CCL3, CCL26, MMP1, and MMP3 (**Fig. 2H**). Notably, IL-1α is a EC-senescence cue that coordinates cytokine signaling networks.^11^ In contrast, canonical pro-inflammatory SASP cytokines such as IL-6, IL-8, and CXCL1 (**Fig. 2I**) were significantly suppressed under stiffer conditions, while IL-11, CCL2, and ICAM-1 levels (**Fig. 2J**) remained unchanged across varying matrix stiffness. This selective suppression of classical SASP factors, alongside enhanced immunomodulatory signals, characterizes a stiffness-responsive secretory profile associated with EC senescence.

Classical SASP, typically triggered by replication stress or DNA damage, is largely mediated by NF-κB (Nuclear Factor-kappa B) signaling,^12,13^ and characterized by robust secretion of IL-6, IL-8, and CXCL1.^14^ By contrast, the SASP induced by matrix stiffening features a distinct cytokine composition, with selective upregulation of IL-33, IL-1α, and IFN-γ molecules known to act upstream of classical SASP mediators.^15-17^ These differences suggest a divergent SASP trajectory under mechanical stress, potentially involving alternative regulatory pathways such as Notch,^18^ MAPK/JNK (Mitogen-Activated Protein Kinase/c-Jun N-terminal kinase)^17^ and JAK/STAT (Janus Kinase/Signal Transducer and Activator of Transcription)^19^.

Mechanical regulation of Notch has been implicated in microvascular remodeling and endothelial phenotypic transitions.^20^ Building on our previous finding that focal adhesion realignment and junctional restructuring mediate mechanotransduction in 3D microvasculatures,^4^ we hypothesized that matrix stiffening induction of Notch ligands drives EC senescence. To test this hypothesis, we examined Notch activation under the matrix stiffening condition. Prior work has linked mechanosensitive Notch to the Notch1-Dll4 axis on softer substrates.^21^ In our 3D model, however, matrix stiffening elevated mRNA for *Notch1* receptor, its ligand *JAG1* and *JAG2*, and the downstream target *HEY1*, with no significant change in *Dll4* ligand transcripts (**Fig. 3A**). Protein analyses also confirmed increased NICD, JAG1, and JAG2 under matrix stiffening (**Fig. 3C-F**). Given the context dependence of Notch receptor-ligand pairing across physiologic and pathologic state,^22-25^ these findings indicate that stiffness cues in a luminalized 3D network preferentially engage a Notch1-JAG axis over Dll4. This mechanically induced shift may reprogram downstream signaling in a stiffness-dependent manner. Consistent with reports implicating Notch1-JAG signaling in pathological vascular remodeling and inflammatory responses,^26^ these results suggest that targeting Notch1 activation could be an effective strategy for stiffness-induced endothelial dysfunction.

**Figure 3:**
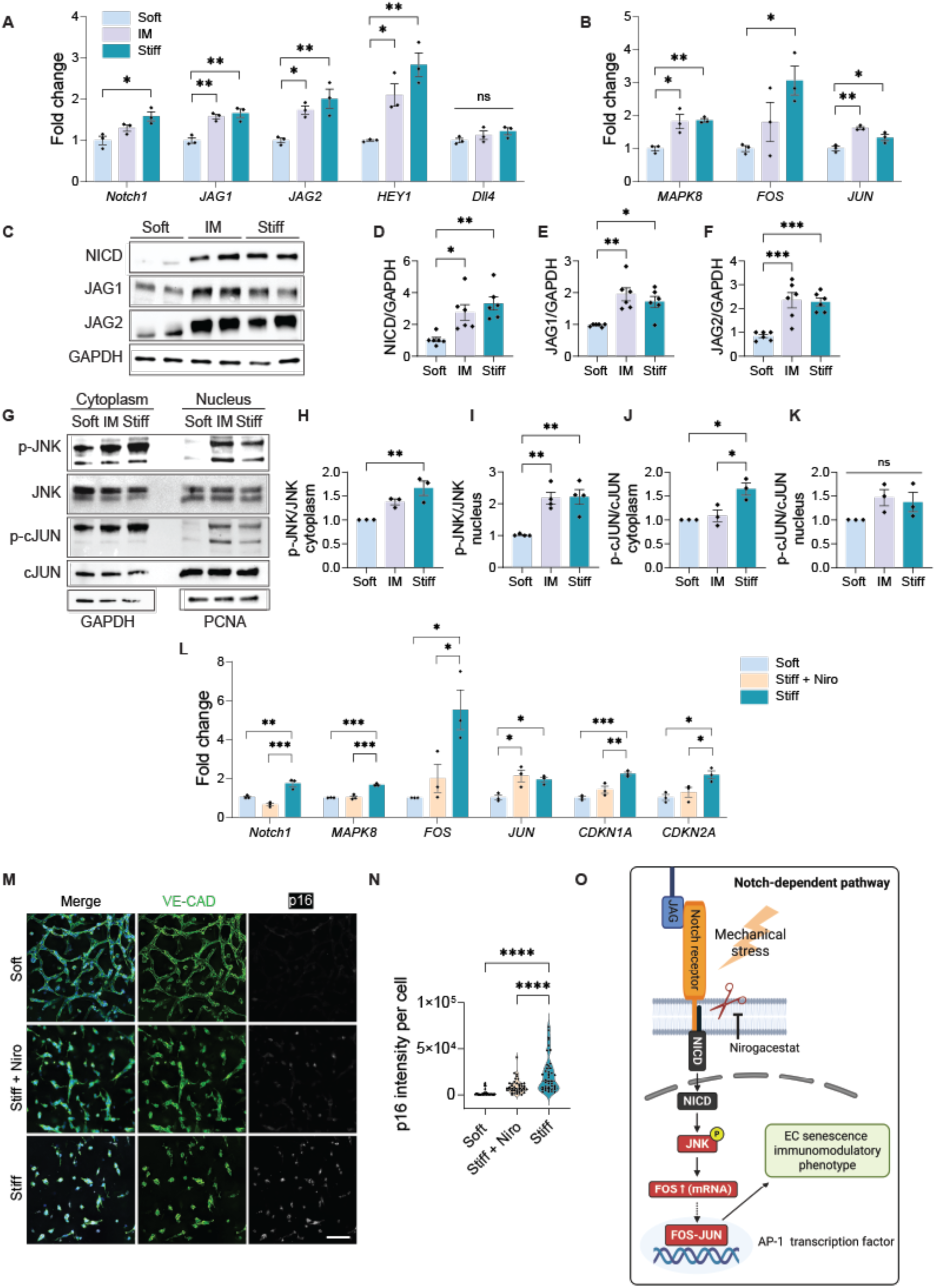
Matrix stiffening activates Notch to drive JNK-FOS signaling and endothelial senescence. (A) qRT-PCR analysis revealed significant upregulation of *Notch1* receptor, ligands *JAG1* and *JAG2*, and downstream target *HEY1* under stiff conditions. *Dll4* ligand gene expression remained unchanged upon matrix stiffening. N=3. (B) Matrix stiffening also induced *MAPK8* (encoding JNK1), *FOS*, and *JUN* gene expression, key components of the JNK-AP-1 signaling axis, as measured by qRT-PCR. N=3. (C-F) Western blot analysis confirmed increased protein levels of cleaved-Notch1 (NICD), JAG1, and JAG2 in stiffened matrices. GAPDH was used as a protein loading control. N=3. (G-K) Matrix stiffening increased phosphorylation of JNK (p-JNK) in both cytoplasmic and nuclear fractions (N=4), while phosphorylation of cJUN (p-c-JUN) was elevated only in the cytoplasm (N=3). This suggest that nuclear p-c-JUN activation is independent of stiffness-mediated Notch signaling. GAPDH and PCNA were used as protein loading controls for cytoplasm and nucleus, respectively. (L) Treatment with nirogacestat (Niro), a γ-secretase inhibitor, reduced stiffness-induced expression of *Notch1*, *MAPK8*, *FOS*, *CDKN1A*, and *CDKN2A* (N=3), indicating that Notch signaling regulates JNK-FOS signaling and senescence-associated cell cycle arrest. (M-N) Immunofluorescence staining exhibited reduced p16 expression in nirogacestat-treated microvascular networks compared to untreated stiff controls. Representative confocal images show endothelial networks stained for VE-Cadherin in green, p16 in white, nuclei in blue. Scale bar is 100 µm. N=3. (O) Schematic illustration of the proposed mechanotransductive pathway: matrix stiffening enhances Notch activation, NICD release, which in turn activates the JNK-FOS axis, promoting endothelial senescence and an immunomodulatory phenotype. Significance levels indicated as ns= not significant (p > 0.05), *p ≤ 0.05, **p ≤ 0.01, and ***p ≤ 0.001.

Beyond its mechanosensitive role, Notch signaling also functions as a molecular scaffold for crosstalk with MAPK/JNK^27,28^, NF-κB^29^, and Wnt (Wingless/Integrated)^30^ pathways, particularly under stress. Accordingly, we examined whether matrix stiffening engages a Notch-dependent JNK-AP-1 module that regulates senescence and inflammation.^25,31,32^ Notably, transcription factor AP-1 induction is required for initiation of stress-induced senescence.^31,33^ At the transcription level, matrix stiffening significantly increased *MAPK8* (encoding JNK1), as well as AP-1 components *FOS* and *JUN* (**Fig. 3B**). Consistently, phosphorylation of JNK (p-JNK) significantly increased in both cytoplasmic and nuclear fractions (**Fig. 3G-I**), indicating JNK pathway activation under mechanical stress.^33^ Whereas prior chemotoxicity studies emphasize nuclear c-JUN phosphorylation (p-c-JUN) during doxorubicin-induced senescence,^33^ in our mechanically driven model, p-c-JUN accumulated in the cytoplasm without a significant nuclear increase (**Fig. 3G, J-K**). Together with robust *FOS* mRNA induction, these data suggest that stiffness-induced senescence proceeds via a FOS-dominant AP-1 program downstream of Notch-JNK, rather than enhanced nuclear c-JUN activity. Importantly, the lack of a significant Notch-dependent increase in nuclear c-JUN activity correlates with the minimal IL-6 and IL-8 secretion observed under matrix stiffening,^33^ implying that incomplete c-JUN activation may restrain development of the classical pro-inflammatory SASP. Notably, time-resolved studies of DNA-damage senescence place JNK/AP-1 activity at early stages of senescence trajectory, and pharmacologic JNK inhibition during this early-to-mid transition has been shown to attenuate subsequent SASP maturation.^33^ In line with this, our 48-h stiffening model, marked by cell cycle arrest, a FOS-skewed AP-1 signature, and limited IL-6/IL-8 secretion, aligns with this early/transitional senescence.^33,34^ By capturing this early window under controlled mechanical cues, our *in vitro* system provides a tractable platform to interrogate upstream regulators before downstream inflammatory circuits become fully engaged.

To place Notch upstream of JNK-AP-1 and assess therapeutic leverage, we treated EC networks with nirogacestat, an FDA-approved γ-secretase inhibitor that blocks Notch receptor cleavage and activation^35^. Given that previous studies have shown that loss of JAG ligands can enhance complementary Notch1–Dll4 signaling,^36^ the use of a pan-Notch inhibitor in our model allowed us to broadly assess the role of Notch signaling in stiffness-induced senescence. Nirogacestat treatment suppressed the stiffness-induced increase in *Notch1*, *MAPK8*, and *FOS* transcripts, while *JUN* mRNA remained elevated (**Fig. 3L**), consistent with c-JUN being maintained by Notch-independent inputs. Functionally, nirogacestat treatment reduced *CDKN1A* and *CDKN2A* gene expression (**Fig. 3L**) and lowered p16 staining intensity compared to untreated Stiff controls (**Fig. 3M-N**), indicating attenuation of stiffness-driven senescence. Together, these findings define a mechanosensitive senescence mechanism that proceeds via a Notch-JNK-FOS axis (**Fig. 3O**), bypassing canonical c-JUN activation and limiting classical SASP. This highlights our model’s capability to uncover early EC senescence programs that expand current understanding of AP-1 function and offer new insights for targeting vascular mechanoaging.

To further validate our *in vitro* findings that matrix stiffness induces EC senescence downstream of Notch signaling in a human model, we examined fibrotic capsule tissue from patients with synthetic breast implants, which reproducibly develop localized fibrosis enriched with senescent cells.^37,38^ Unlike chronic fibrotic diseases characterized by prolonged systemic inflammation and complex immune pathologies, this foreign body response model reflects localized fibrotic remodeling driven predominantly by mechanical forces and the local microenvironment.^39^ Given that our *in vitro* hydrogel platform recapitulates a mechanically induced EC senescence program without chronic, systemic inflammatory confounders, we leveraged this model as a relevant *in vivo* system to study mechanically driven fibrosis and senescence. To assess EC senescence and Notch signaling *in vivo*, we performed triple immunofluorescence staining for CD31 (endothelial marker), p16 (senescence marker), and Notch1 (Notch signaling marker) in both fibrotic and adjacent soft control tissues (muscle and fat). The proportion of CD31^+^p16^+^ double-positive cells was significantly higher in fibrotic capsules, confirming the accumulation of senescent ECs in stiffened microenvironments (**Fig. 4A-B**). While the percentage of CD31^+^Notch1^+^ double-positive cells, representing Notch1 expression in non-senescent ECs, remained similar between soft and fibrotic tissues (**Fig. 4C**), the fraction of triple-positive (CD31^+^p16^+^Notch1^+^) cells was significantly higher in fibrotic capsules (**Fig. 4D**). This indicates that Notch1 receptor expression is preferentially increased in senescent ECs *in vivo* under fibrotic stiffening. Quantification of vessel area distribution showed no significant differences between soft control and fibrotic tissues (**Fig. S3**).

**Figure 4:**
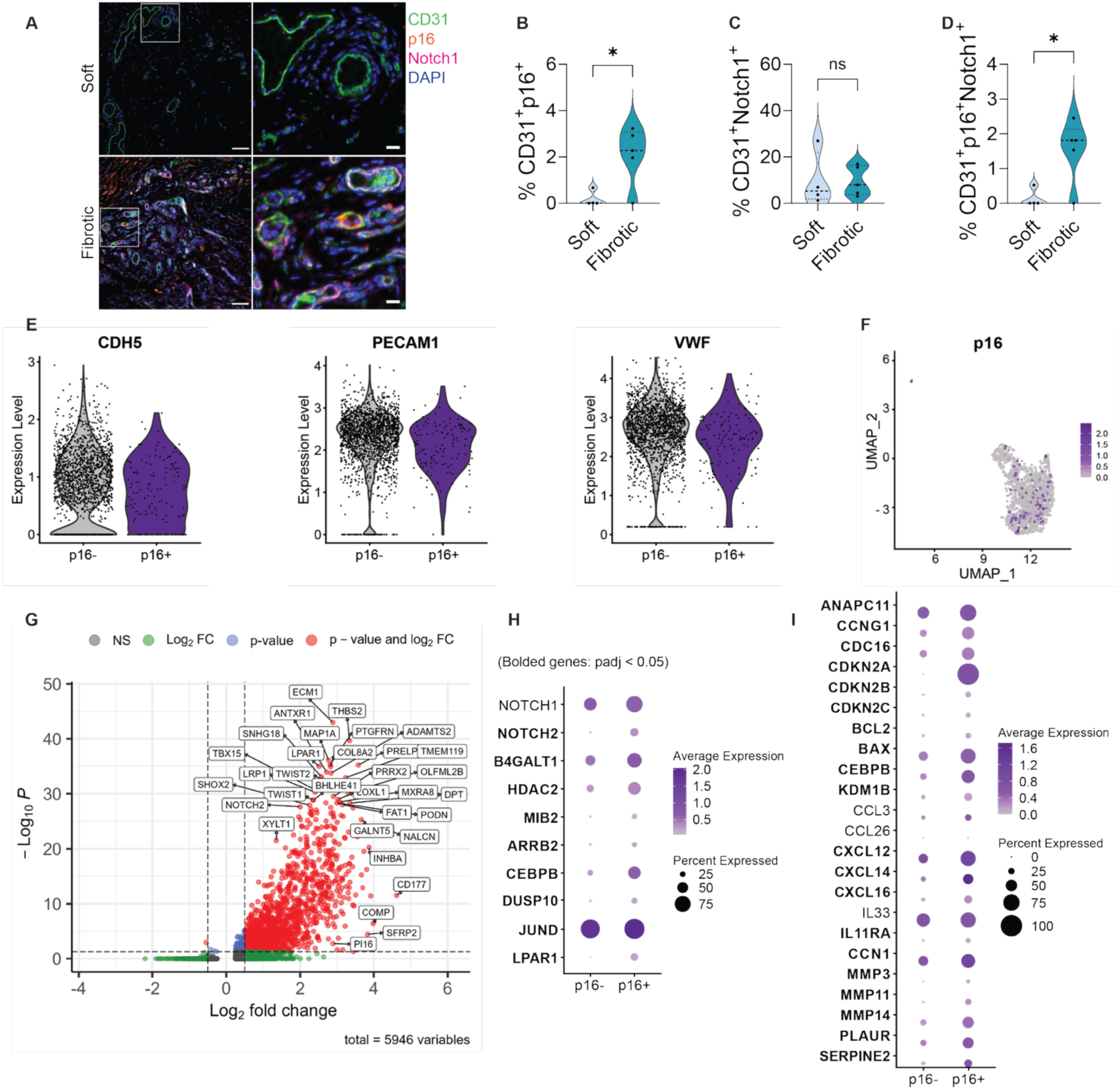
Human breast implant fibrotic capsule tissue confirms Notch-driven endothelial senescence under stiffened environment. (A) Immunofluorescence staining of fibrotic capsules surrounding surgically explanted synthetic breast implants and adjacent fat tissue showed ECs (CD31) in green, senescence marker (p16^INK4a^) in orange, Notch1 receptor in magenta, nuclei in blue. Representative images of both soft and fibrotic tissues were obtained from the same patient. Scale bar = 100 μm (left), 20 μm (right). (B) Increased p16 expression was observed in ECs within fibrotic capsule regions compared to soft control tissues, including fat and muscle. (C) The proportion of non-senescent ECs expressing Notch1 (CD31^+^Notch1^+^) was comparable between soft control and fibrotic tissues, suggesting Notch1 expression in non-senescent ECs is not altered by stiffness. (D) A significantly higher frequency of triple-positive (CD31^+^p16^+^Notch1^+^) cells in fibrotic regions indicates that Notch 1 expression is enriched specifically within senescent ECs in stiffened environments. (E) Violin plots confirm EC identity through expression of standard EC markers VE-Cadherin (CDH5), PECAM1, and von Willebrand factor (vWF). (F) UMAP projection identifies 132 p16⁺ ECs out of 1,836 total ECs. (G) Volcano plot showing differentially expressed genes between p16⁺ and p16⁻ ECs. Genes meeting adjusted p < 0.05 and log₂ fold change > 0.5 are highlighted. (H) Dot plot displays enrichment of Notch- and JNK-associated genes in senescent ECs. (I) p16⁺ ECs show increased expression of genes related to cell cycle arrest (e.g., CDKN2A/B/C, CCNG1), SASP and inflammatory signaling (e.g., BCL2, BAX, CXCL12/14/16, IL11RA), and ECM remodeling (e.g., MMP3/11/14, PLAUR, SERPINE2, CCN1), supporting their active role in fibrosis-associated tissue remodeling.

We next analyzed the endothelial subset from a previously published single-cell RNA sequencing dataset of human fibrotic capsules surrounding synthetic breast implants.^37^ Senescent ECs were identified based on expression of the cell cycle inhibitor *CDKN2A* (which encodes *16^Ink4a^)*, alongside standard EC markers, including *Cadherin-5* (*VE-CAM*), *PECAM-1*, and *vWF* (**Fig. 4E**).^40,41^ Across the Uniform Manifold Approximation and Projection (UMAP) space, 132 of the total 1,836 ECs (∼7%) expressed p16 (**Fig. 4F**), consistent with the expected low frequency of senescent cells *in vivo*.^37^ This proportion is comparable to our immunofluorescence-based quantification, which identified ∼12% of CD31^+^ cells were p16^+^, supporting the consistency of senescent EC detection across experimental modalities.

Differential gene expression analysis between p16^+^ and p16^-^ ECs identified 1487 genes with significant changes (adjusted p < 0.05). These genes were enriched in functional categories related to inflammatory signaling, ECM remodeling, and chromatin regulation (**Fig. 4G**). While the volcano plot displays all genes detected in the analysis, only those meeting both statistical significance (adjusted p < 0.05) and a log₂ fold change > 0.5 are highlighted as robustly regulated targets.

We next asked whether the transcriptomic features of p16^+^ ECs reflected the Notch-JNK senescence axis observed *in vitro*. Analysis showed that p16^+^ ECs expressed several key components of Notch pathway components (*Notch1/2, B4GALT1, HDAC2,* and *MIB2*) and JNK-associated components (*ARRB2*, *CEBPB*, *DUSP10*, *JUND*, and *LPAR1*) (**Fig. 4H**). Although Notch1 was expressed in a greater proportion of p16^+^ cells, its expression level was comparable between the p16^+^ and p16^-^ groups.

Senescence-associated genes were further profiled (**Fig. 4I**). Genes involved in cell cycle regulation (*ANAPC11*, *CCNG1*, *CDC16*, *CDKN2A*, *CDKN2B*, and *CDKN2C*) were differentially expressed in p16^+^ cells, consistent with a senescent phenotype. SASP-related genes, including *BCL2* and *BAX* genes, as well as chromatin regulators such as *CEBPB* and *KDM1B*, were also enriched in the p16^+^ population. Moreover, immune signaling molecules such as *CXCL12*, *CXCL14*, *CXCL16*, and *IL11RA* showed elevated expression, suggesting an immunomodulatory and pro-inflammatory profile in senescent ECs. Importantly, genes identified from our *in vitro* Luminex analysis, including *CCL3*, *CCL26*, and *IL33*, were expressed by a higher proportion of p16⁺ cells, even though their average expression levels remained similar between groups.

Lastly, ECM remodeling genes such as *CCN1*, *MMP3*, *MMP11*, *MMP14*, *PLAUR*, and *SERPINE2* were significantly upregulated in the p16^+^ group (**Fig. 4I**). Gene Set Enrichment Analysis (GSEA) further confirmed enrichment of pathways related to collagen fibril organization and collagen metabolism in the senescent EC population (**Fig. S4**). While ECs are not the primary producers of fibrotic matrix, these findings imply that senescent ECs may contribute to local ECM remodeling and potentially facilitate the progression of fibrosis. This highlights the broader impact of EC senescence in shaping tissue remodeling and the development of fibrotic microenvironments.

**Fig S3.**
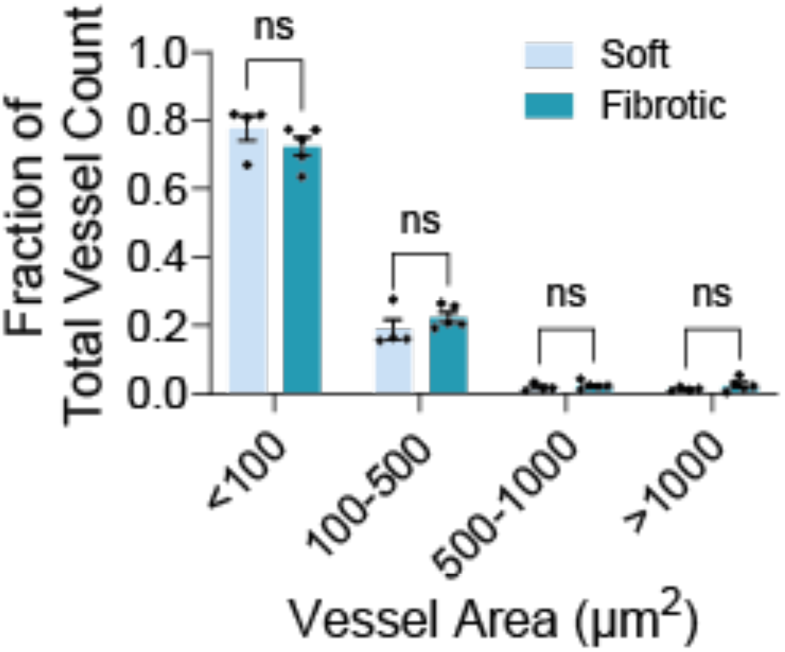
Quantification of endothelial vessel area distribution in soft vs. fibrotic breast capsule tissues. Bar graph showing the fraction of the total vessel count across four vessel area categories. Vessels were quantified from immunofluorescence image based on CD31 staining. No significant differences were observed between groups.

**Fig S4.**
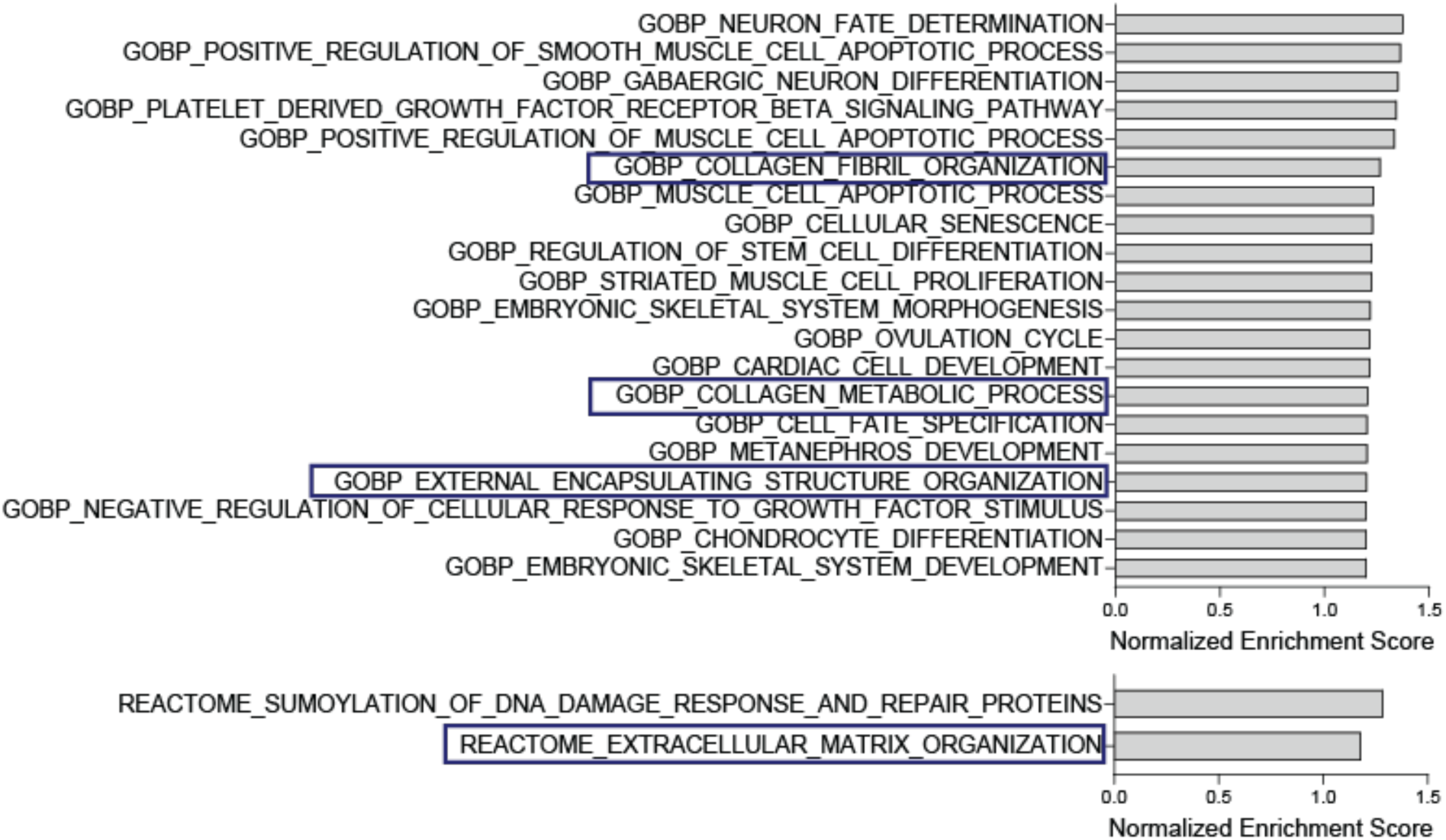
Gene Set Enrichment Analysis (GSEA) of p16^+^ ECs showing enrichment of pathways (adjusted p < 0.05) related to collagen fibril organization, collagen metabolism, and extracellular organization.

Together, our findings establish matrix stiffness as a key upstream driver of EC senescence through a Notch-JNK-FOS signaling axis that promotes a FOS-dominant AP-1 program over canonical c-JUN-driven responses. By capturing a transitional senescence state characterized by a non-classical, immunomodulatory SASP, our hydrogel platform reveals a mechanosensitive senescent phenotype in ECs. Integration with an *in vivo* human fibrotic capsule model further validates the presence of Notch1-expressing ECs in stiffened environments and highlights their potential role in modulating ECM dynamics. This work not only positions our system as a novel platform for dissecting early, targetable stages of mechanoaging, but further expands our understanding of how mechanical cues shape vascular aging through non-canonical molecular trajectories.

## Experimental Section

### Synthesis of Methacrylated HA (HA–MA)

HA-MA was synthesized as in previously reported.^42^ Approximately 14 or 64% of the primary hydroxyl groups modified with methacrylate groups. Briefly, 2% (w/v) solution of HA (Lifecore, 60 kDa) was prepared on ice and reacted with methacrylic anhydride (Sigma) at HA/MA molar ratios of 1: 0.5 and 1:3 to achieve ∼14% and ∼64% modification, respectively. The pH was maintained at 8–9 for 4 h. The products were dialyzed against NaCl Solution for 48 h, lyophilized, and the degree of modification was determined with ^1^H NMR (Bruker).

### Formation and Characterization of Hydrogels

Stock solutions of Col-MA (8 mg/mL, Advanced Biomatrix) and HA-MA (20 mg/mL in PBS) were prepared separately. To initiate collagen physical crosslinking, a defined volume of neutralization solution (Advanced Biomatrix) was added to the Col-MA solution to adjust pH. The pH-adjusted solution was then mixed with HA-MA solution and PBS to achieve final concentration of 2.5 mg/mL Col-MA and 0.25 mg/mL HA-MA. The resulting precursor mixture was incubated at 37 °C for 30 min to form a physically crosslinked Col-HA hydrogel. Following gelation, the photoinitiator was prepared according to the manufacturer’s instruction (Advanced Biomatrix). Briefly, Ruthenium (Ru, 28 mg/mL) and Sodium Persulfate (SPS, 119 mg/mL) were diluted in PBS to final concentrations of 0.7 mg/mL and 2.4 mg/mL, respectively. This solution was added on top of the gels and incubated for 10 min to allow diffusion into the gels. To induce secondary chemical crosslinking and stiffening, hydrogels were then exposed to white LED light for 1 min.

Hydrogel bulk stiffness was characterized using a Discovery Hybrid Rheometer (HR20, TA Instruments) equipped with an 8 mm cross-hatched geometry and a 500 um gap, maintained at 37 °C. Premade hydrogel disks (140 μL, 8 mm diameter) were placed on the rheometer stage, and PBS was added around the geometry to ensure the hydrogels remained swollen. An amplitude sweep was performed from 0.1% to 10% strain at a constant frequency of 0.25 Hz. Storage modulus (G’) values were recorded at 0.1% strain within the linear viscoelastic regime. Young’s modulus (E) was calculated from the shear modulus using the equation: E = 2G(1+ν) (E is Young’s modulus, G is shear modulus and ν is Poisson’s ratio), assuming a Poisson ratio of 0.5^43^.

To assess the equilibrium swelling ratio, premade hydrogels were formed in pre-weighted cell culture insert wells. After full swelling, the weight of the hydrated hydrogels were measured. The dry hydrogel weight was obtained by incubating the hydrogels at 60 °C until completely dehydrated. The swelling ratio (%) was calculated as:

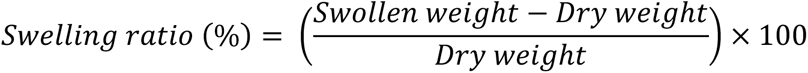

Collagen-HA fibers were imaged using reflective microscopy (640 nm, Nikon Eclipse Ti2 microscope). Fiber diameter and coverage area were quantified using Nikon NIS software. A total of 25 images acquired from 4 replicates were analyzed for quantification.

### Cell Culture

All cells were maintained in a humidified incubator at 37 °C with 5% CO_2_. ECFCs (provided by the Yoder Lab, Indiana University School of Medicine) were cultured in EGM2 (Lonza) supplemented with an additional 10% HyClone FBS, beyond the standard formulation. For 2D expansion, cells were seeded on type I collagen (Corning)-coated tissue culture plates. For 3D hydrogel cultures, the media was further supplemented with 50 ng/mL of VEGF (PeproTech) and 25 ng/mL of bFGF (PeproTech) to support endothelial network formation.

### 3D Cell Encapsulation and Dynamic Stiffening

The hydrogel precursor mixture was prepared and neutralized as described above. ECFC were suspended in EGM2 supplemented with 50 ng/mL VEGF and 25 ng/mL bFGF, then mixed with the hydrogel solution to achieve a final cell concentration of 2 million cells/mL. Dynamic stiffening was applied 48 h after cell encapsulation by exposing the cell-laden hydrogel constructs to LED light, as described above.

### Hydrogel Immunofluorescent staining, Imaging, and Quantification

On day 4, cell-laden hydrogel constructs were fixed with 4% (w/v) paraformaldehyde in PBS for 45 min and then blocked/permeabilized with 5% (w/v) bovine serum albumin and 0.1% (v/v) Triton X-100 for 4 h at room temperature. The constructs were incubated with primary antibodies, including anti rabbit-p16 (1:400, Proteintech) and anti mouse-VE-CAD (1:400, Santa Cruz) overnight at 4 °C. Subsequently, the constructs were incubated with corresponding secondary antibodies or phalloidin (Invitrogen). Nuclei were stained with DAPI. Fluorescent images were taken by Nikon EclipseTi2 microscope using Z-stack mode. Maximum intensity projection images were generated by Nikon NIS analysis software. Image quantification was conducted using ImageJ and Imaris.

SA-β-Gal staining was performed using CellEvent™ Senescence Green Detection Kit (Thermo Fisher) according to the manufacturer’s instructions with minor modifications. Day4 cell-laden hydrogel constructs were fixed and then blocked/permeabilized with 1% (w/v) bovine serum albumin and 0.1% (v/v) Triton X-100 for 30 min at room temperature. The constructs were subsequently incubated with prewarmed working solution for 3 h at 37 °C without CO_2_ and protected from light. Phalloidin (Invitrogen) and DAPI were used for counterstaining. Imaging and quantification were performed as described above.

### Luminex Analysis

Custom Luminex assay kits (Bio-Techne) were used to quantify soluble factors. Cell culture supernatants were collected on day 4 of culture from hydrogels under Soft, IM, and Stiff conditions (N=2, n=5). Fresh cell culture media served as blank control. All Luminex assays were performed in the Regional Biocontainment Laboratory (RBL) at Duke University, following the manufacturers’ instructions. Analyte concentrations were normalized to dsDNA content, measured using the PicoGreen assay (Invitrogen, N=2, n=7), following previously published procedures^44^.

### Small-Molecule Inhibition Studies

Nirogacestat, a γ-secretase inhibitor, was used to block Notch receptor cleavage and activation. A 10 mM stock solution of nirogacestat in DMSO was diluted to a final concentration of 750 nM in EGM2 supplemented with 50 ng/mL VEGF and 25 ng/mL bFGF. The inhibitor-containing media was added to cell-laden hydrogel constructs 3 hours prior to matrix stiffening (day 2 post-encapsulation). Nirogacestat treatment was maintained during the stiffening process and continued through day 4 of culture.

### Gene Expression

After 4 days of culture, cell-laden constructs were flash-frozen in liquid nitrogen and stored at −80 °C until further processing. Frozen samples were thoroughly homogenized in TRIzol (Invitrogen) and extracted RNA was purified using the RNA Clean &Concentrator-5 Kit (Zymo Research) according to the manufacturer’s protocol. Complimentary DNA was synthesized using GoScript Reverse Transcription Mix (Promega). Quantitative real-time PCR was performed using Maxima SYBR Green/Fluorescein Master Mix (ThermoFisher). Primer sequences are listed in *Table S1*. The *MAPK8* and *FOS* primers were purchased from Millipore Sigma. GAPDH was used as a reference gene. The fold change was processed by qbase+ software (Biogazelle). A total of three biological repeats were analyzed for each gel composition.

**Table S2.**
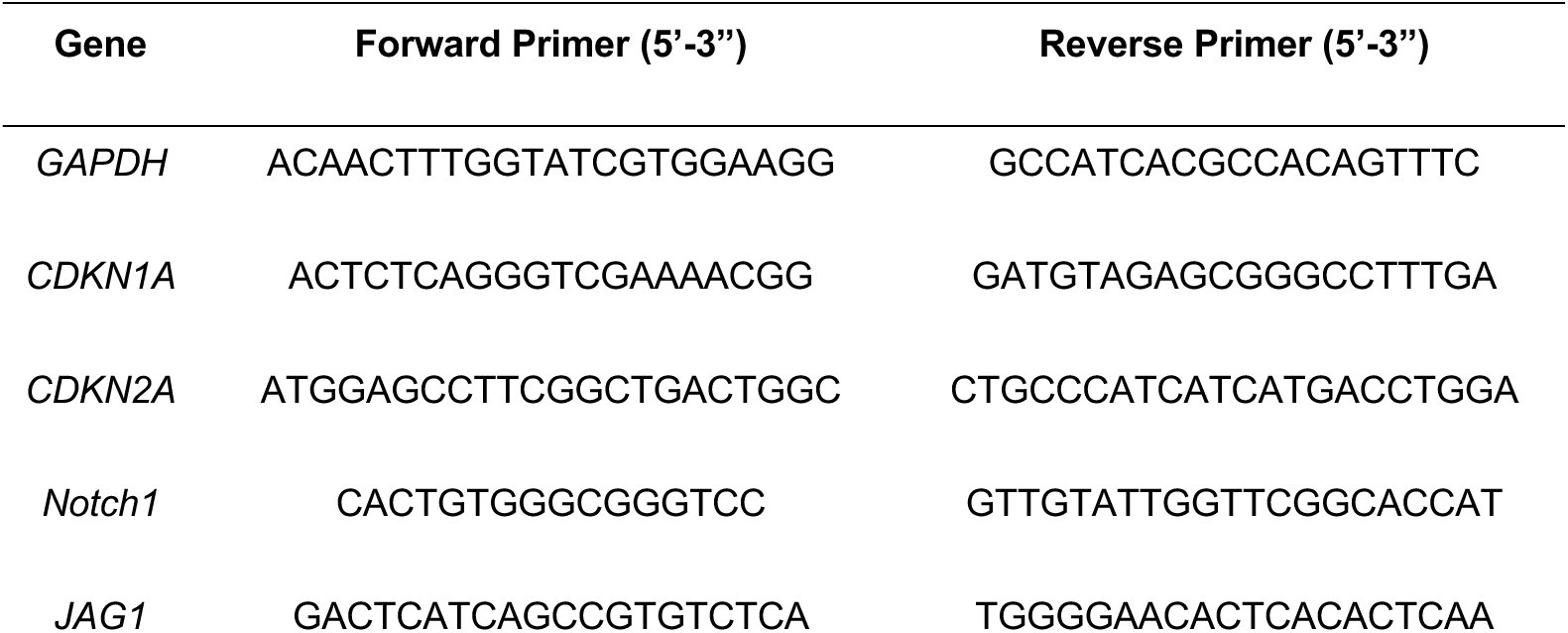
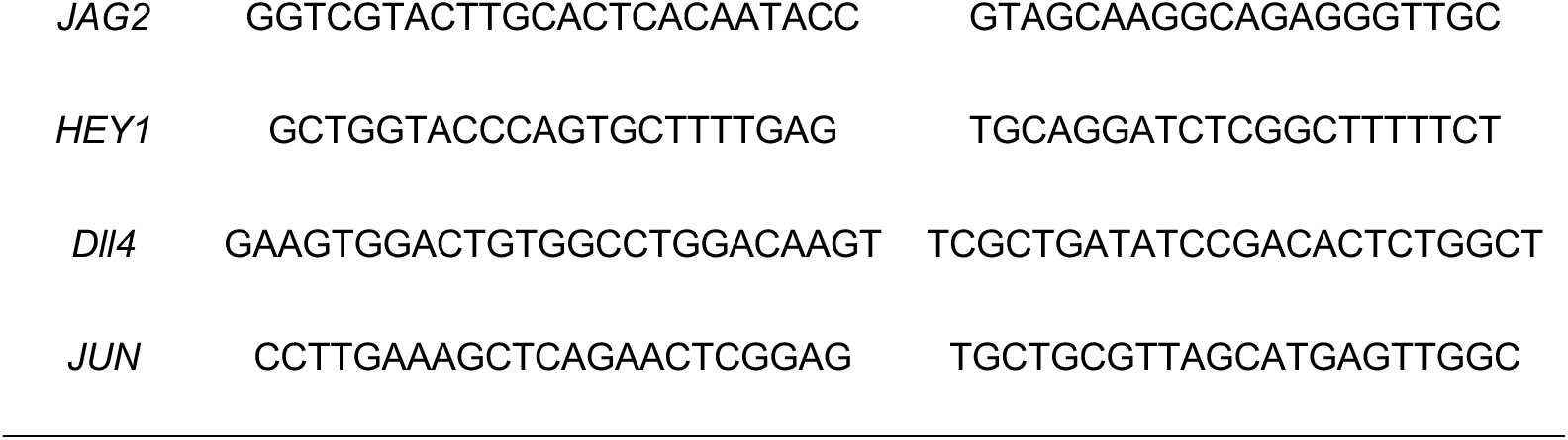
Forward and reverse primers used for qPCR analysis.

### Western Blotting

Day 4 frozen constructs were lysed in RIPA buffer (Thermo Fisher) containing 1% (v/v) of Halt protease and phosphatase inhibitor cocktail (Thermo Fisher). For nuclear and cytoplasmic protein separation, the NE-PER Nuclear and Cytoplasmic Extraction Reagent kit was used following the manufacturer’s instructions. Protein concentrations were determined using the BCA assay kit (Thermo Fisher). Samples were denatured by boiling at 95 °C for 10 min, and 20−35 μg of protein was loaded per lane on 4–20% or 7.5% precast polyacrylamide gel (Bio-Rad). Proteins were transferred to Immun-Blot PVDF Membrane (Bio-Rad) overnight at 4 °C. Protein transfer was confirmed by Ponceau-S staining. Membranes were blocked for 1 h with 5% BSA in TBST for detection of phosphorylated proteins, and 5% milk in TBST for non-phosphorylated proteins. Membranes were then incubated overnight at 4 °C with primary antibodies targeting: p21 (Cell Signaling, rabbit, 1:1000), NICD (Cell Signaling, rabbit, 1:1000), JAG1 (Cell Signaling, rabbit, 1:1000), JAG2 (Cell Signaling, rabbit, 1:1000), phospho-JNK (Cell Signaling, rabbit, 1:750), JNK (Cell Signaling, rabbit, 1:750), phospho-cJUN (Cell Signaling, rabbit, 1:750), cJUN (Cell Signaling, rabbit, 1:750), and PCNA (Proteintech, rabbit, 1:1000). Following primary antibody incubation, blots were washed three times with TBST and incubated with species-appropriate HRP-conjugated secondary antibodies or HRP-conjugated GAPDH (Cell Signaling, rabbit, 1:2500). Immunoblots were detected by an enhanced chemiluminescence western blotting substrate (Thermo Fisher) and imaged using ChemiDoc ImagingSystem (Bio-Rad). Quantification was performed using Bio-Rad Image Lab software. Phosphorylated protein levels were normalized to their total protein counterparts. Original blots were included in Fig. S5.

**Fig S5.**
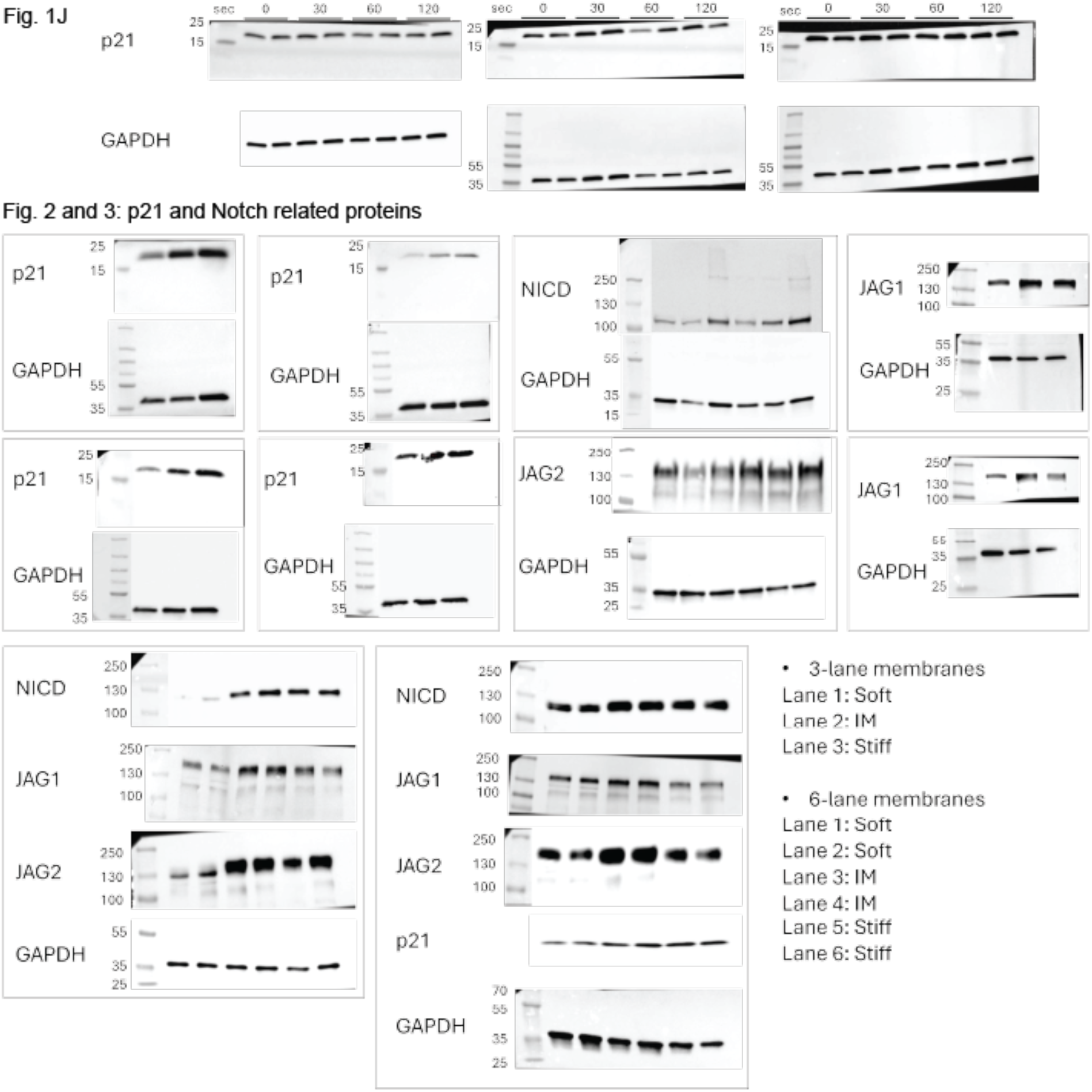
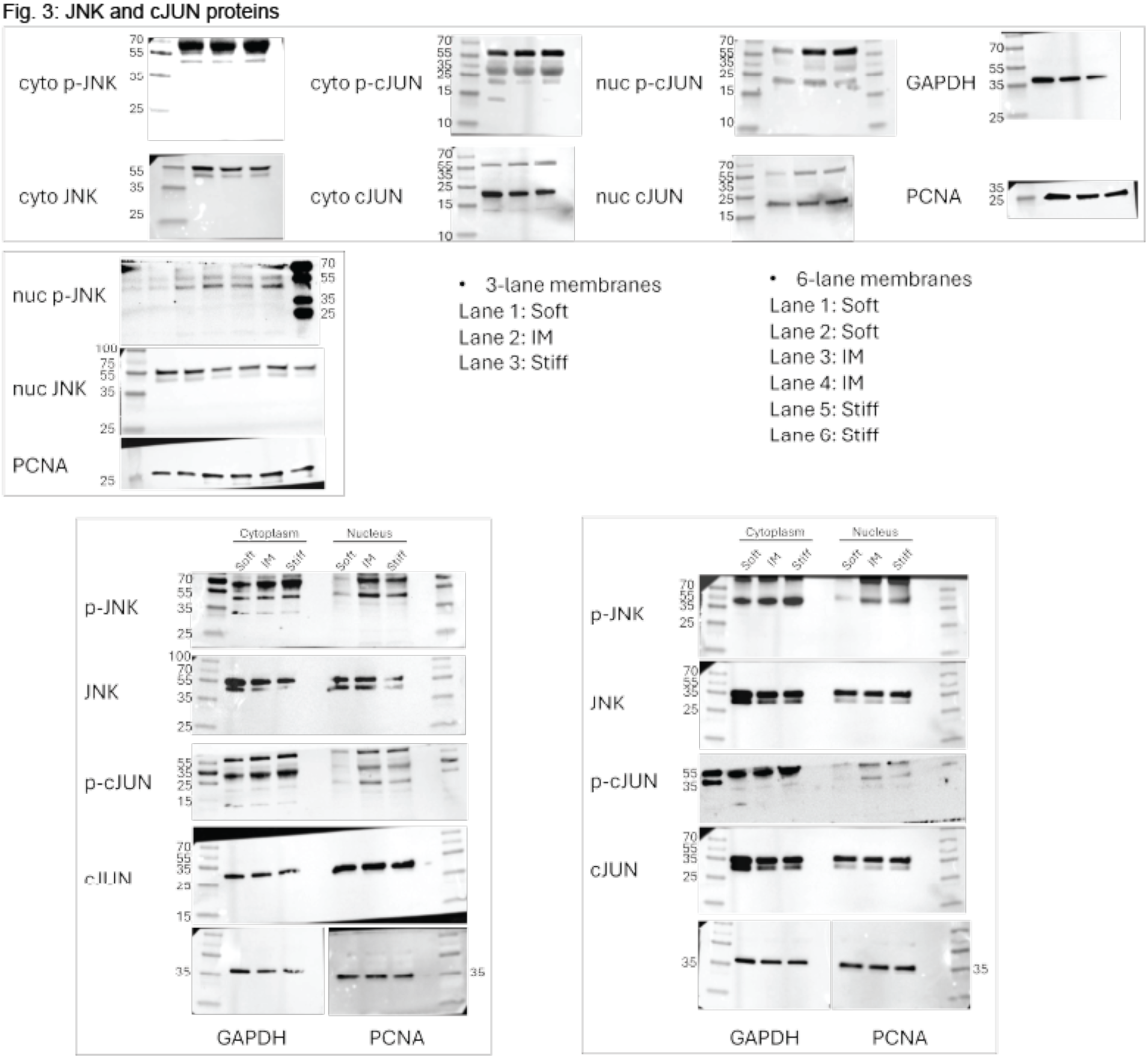
Western blot membranes corresponding to the main figures are shown.

### Human Samples

Deidentified surgical discards from patients undergoing breast implant exchange or replacement surgeries were collected under Johns Hopkins University Institutional Review Board Exemption IRB00088842. For single-cell RNA-sequencing, cells were isolated from the fibrotic capsules from one patient, as previously published.^37^ For immunofluorescence staining and imaging, discarded fibrotic capsule tissue and implant-adjacent muscle and fat tissue were obtained from six patients. All tissues were immediately fixed in 10% formalin for 48 hours. Tissues were then rinsed twice in PBS, dehydrated using a graded ethanol series (70%, 80%, 95%, 100%, 100% ethanol for 1 hour each), and cleared in xylene for 1-2 hours. Tissues were incubated in 4-6 exchanges of melted paraffin wax at 60°C for 24 hours to prepare for embedding. Tissues were embedded into wax blocks using metal molds and stored at room temperature until sectioning. Embedded tissues were sectioned at 5 μm using a microtome.

### Single Cell RNA Sequencing

Single cell RNA-sequencing data were obtained from a previously published dataset of cells isolated from human breast implant fibrotic capsule tissues.^37^ All analyses were performed using the Seurat v5.1 package.^45^ Endothelial cells were identified based on their expression of known markers (PECAM1, CDH5, VWF) and were subsetted for downstream analysis. Cells that were positive for CDKN2A (p16) expression were annotated as senescent. All p16+ cells were negative for proliferation marker MKI67. Differential expression analysis was performed between senescent (p16+) and non-senescent (p16-) cells using a Wilcoxon Rank Sum test via the FindMarkers function in Seurat, with thresholds of logFC≥0.25 and min.pct=0.1. Genes with an adjusted p-value below 0.05 were considered differentially expressed. Differentially expressed genes were visualized using the Seurat, EnhancedVolcano v1.22.0^46^, and ggplot2 packages. Gene set enrichment analysis (GSEA) was performed using the FindMarkers function in Seurat followed by the Fast Gene Set Enrichment Analysis (fgsea) v1.30.0 package^47^. Genes were ranked for GSEA analysis using two steps. First, differential expression was performed using the FindMarkers function with thresholds of logFC≥0, min.pct=0, minimum.cells.group=1, and min.cells.feature=1. Then, genes were ranked in descending order by their values of the following calculation: -log10(p-value) * avg_log2FC. Ranked genes were input into fgsea to identify pathways enriched in senescent endothelial cells. GSEA was performed using the human HALLMARK, REACTOME, KEGG, and GOPB databases. Pathways with a false discovery rate of >0.05 were considered significant.

### Immunofluorescence Staining of Human Tissues

Tissue sections were stained using the PhenoCode Signature platform by Akoya Biosciences. Slides were incubated in an oven at 60°C to remove excess paraffin wax. Then, slides were deparaffinized in xylenes for 3x3 min and rehydrated in a graded series of ethanol (100% x 2, 95%, 80%, 70%; 1 min each) and Millipore water (3x3 min). Antigen retrieval was performed by incubating slides in pre-heated 1X AR6 buffer in a vegetable steamer for 15 min. Slides were cooled at RT for 15 min, rinsed in Millipore water, and incubated in 3% H_2_O_2_ for 15 min to block endogenous peroxidases. Then, slides were rinsed once in Millipore water and incubated in a blocking buffer of 10% bovine serum albumin and 0.05% Tween-20 in 1X PBS for 30 min at RT. Slides were stained with the following antibodies in blocking buffer at RT: p16 for 15 min (CINTEC-Roche, 705-4793, 1:5), Notch1 for 30 min (Cell Signaling, 3608, 1:200), and CD31 for 30 min (Abcam, ab76533, 1:100). Following the primary antibody incubation, slides were washed in 1X TBST for 3x3 min and incubated in secondary antibody for 30 min at RT. We used MACH 4 Universal HRP-Polymer (Biocare Medical, M4U534) to detect p16 and MACH 3 Rabbit HRP Polymer Detection (Biocare Medical, M3R531) to detect Notch1 and CD31. After TBST washes, slides were incubated in Opal dyes diluted in 1X Manual Amplification Diluent for 10 min at RT. Opal dyes 570, 650, and 520 were used to label p16, Notch1, and CD31, respectively. Slides were washed in TBST for 3x3 min. Incubations with AR6 buffer, blocking buffer, primary antibodies, secondary antibodies, and Opal dyes were repeated three times to enable co-staining of p16, Notch1, and CD31. Only one antibody was used for staining in each round. Following the three rounds of antibody staining, slides were rinsed in Millipore water, stained with 1X Spectral DAPI solution for 5 min, rinsed twice in Millipore water, and mounted in DAKO Fluorescence Mounting Medium (Agilent, S302380-2) using #1.5 coverslips. Slides were dried overnight and stored at 4°C until imaging. Slides were imaged using the Zeiss Axio Observer.Z1/7 with Apotome and processed using ZenBlue software.

### Colocalization and Vessel Area Analysis of Human Tissues

Images of human breast implant tissues were analyzed for colocalization of senescence and vascular markers (CD31, Notch1, and p16) using HALO v3.6 with the HighPlex FL v3.2 module (Indica Labs). Regions of interest (ROIs) were drawn to select the regions with the fibrotic capsules (stiff) and muscle or fat (soft) in each image. Nuclear detection was performed using the DAPI stain and cytoplasmic detection was used for the antibody stains. Minimum intensity thresholds were set for each stain and were kept consistent across images. Primary delete slides for each antibody stain were used as negative controls to minimize false positive signal detection. Colocalization of the antibody stains in each DAPI+ cell were measured, exported, and analyzed in GraphPad Prism 10.

### Vessel area and density measurements were performed using a custom script in QuPath

0.5.1. ROIs were drawn using the same approach as in the colocalization analysis. Vessel detection was performed using the OpenCVPixelClassifer function. A gaussian blur and minimum intensity threshold were applied to detect areas positive for CD31 (vessel marker). Vessel areas <20 μm^2^ or >900,000,000 μm^2^ were excluded from the analysis. The resulting vessel annotation in each image was split to enable the analysis of individual vessel fragments. Tissue ROI and vessel area measurements were exported and analyzed using GraphPad Prism 10.

### Statistical Analysis

The authors performed statistical analysis using GraphPad Prism 10 (GraphPad Software Inc.). This software was also used to perform t-tests and one-way ANOVA to determine significance. Replicates were indicated throughout the figure captions. Significance levels were set at *p < 0.05, **p < 0.01, ***p < 0.001, and ****p < 0.0001.

## Acknowledgements

Luminex assay was performed in the Regional Biocontainment Laboratory (RBL) at Duke University, which received partial support for construction and renovation from the National Institutes of Health (UC6-AI058607 and G20-AI167200), and facility support from the National Institutes of Health (UC7-AI180254). Tissue sectioning was conducted by the Johns Hopkins University Oncology Tissue and Imaging Service (OTIS) Core Laboratory (NIH P30 CA006973). HALO image analysis was conducted using resources provided by the Johns Hopkins University Tumor Microenvironment (TME) Core.

We thank Nicole Hanson for assistance with western blotting, Makenzie Bushold for support with immunofluorescence staining, and Yaqing Huang for helpful discussion regarding single-cell RNA sequencing. We also thank Yusef Mathkour for his advice on the vessel image analysis.

This work was supported in part by American Heart Association (AHA, 25POST1359525), Mandel Foundation, Duke Regeneration Center (DRC), and Duke Science and Technology. Additional funding for the human sample study was provided by NIH (U54AG079779, R01AG082965) and the Bloomberg ∼ Kimmel Institute.

## References

1 Selman, M. & Pardo, A. Fibroageing: An ageing pathological feature driven by dysregulated extracellular matrix-cell mechanobiology. Ageing Research Reviews 70, 101393 (2021).

2 Lu, T. & Finkel, T. Free radicals and senescence. Experimental cell research 314, 1918–1922 (2008).

3 Wang, D., Brady, T., Santhanam, L. & Gerecht, S. The extracellular matrix mechanics in the vasculature. Nat Cardiovasc Res 2, 718–732 (2023). 10.1038/s44161-023-00311-0

4 Schnellmann, R. et al. StiUening matrix induces age-mediated microvascular phenotype through increased cell contractility and destabilization of adherens junctions. Advanced Science 9, 2201483 (2022).

5 Brandon, K. D., Frank, W. E. & Stroka, K. M. Junctions at the crossroads: the impact of mechanical cues on endothelial cell-cell junction conformations and vascular permeability. Am J Physiol Cell Physiol 327, C1073–C1086 (2024). 10.1152/ajpcell.00605.2023

6 Bloom, S. I., Islam, M. T., Lesniewski, L. A. & Donato, A. J. Mechanisms and consequences of endothelial cell senescence. Nature Reviews Cardiology 20, 38–51 (2023).

7 Wang, E. Y., Zhao, Y., Okhovatian, S., Smith, J. B. & Radisic, M. Intersection of stem cell biology and engineering towards next generation in vitro models of human fibrosis. Front Bioeng Biotechnol 10, 1005051 (2022). 10.3389/fbioe.2022.1005051

8 Ahmed, D. W. et al. Integrating mechanical cues with engineered platforms to explore cardiopulmonary development and disease. iScience 26, 108472 (2023). 10.1016/j.isci.2023.108472

9 Manon-Jensen, T., Kjeld, N. & Karsdal, M. A. Collagen-mediated hemostasis. Journal of Thrombosis and Haemostasis 14, 438–448 (2016).

10 Rhodes, J. M. & Simons, M. The extracellular matrix and blood vessel formation: not just a scaUold. Journal of cellular and molecular medicine 11, 176–205 (2007).

11 Barinda, A. J. et al. Endothelial progeria induces adipose tissue senescence and impairs insulin sensitivity through senescence associated secretory phenotype. Nature communications 11, 481 (2020).

12 Chien, Y. et al. Control of the senescence-associated secretory phenotype by NF-κB promotes senescence and enhances chemosensitivity. Genes & development 25, 2125–2136 (2011).

13 Wang, B., Han, J., ElisseeU, J. H. & Demaria, M. The senescence-associated secretory phenotype and its physiological and pathological implications. Nature Reviews Molecular Cell Biology 25, 958–978 (2024).

14 Moiseeva, V. et al. Senescence atlas reveals an aged-like inflamed niche that blunts muscle regeneration. Nature 613, 169–178 (2023).

15 Orjalo, A. V., Bhaumik, D., Gengler, B. K., Scott, G. K. & Campisi, J. Cell surface-bound IL-1α is an upstream regulator of the senescence-associated IL-6/IL-8 cytokine network. Proceedings of the National Academy of Sciences 106, 17031–17036 (2009).

16 Chan, B. C., Lam, C. W., Tam, L.-S. & Wong, C. K. IL33: roles in allergic inflammation and therapeutic perspectives. Frontiers in immunology 10, 364 (2019).

17 Chow, J., Wong, C. K., Cheung, P. F. & Lam, C. W. Intracellular signaling mechanisms regulating the activation of human eosinophils by the novel Th2 cytokine IL-33: implications for allergic inflammation. Cellular & molecular immunology 7, 26–34 (2010).

18 Hoare, M. et al. NOTCH1 mediates a switch between two distinct secretomes during senescence. Nature cell biology 18, 979–992 (2016).

19 Pan, J. et al. Mechanical stretch activates the JAK/STAT pathway in rat cardiomyocytes. Circulation research 84, 1127–1136 (1999).

20 Mack, J. J. & Iruela-Arispe, M. L. NOTCH regulation of the endothelial cell phenotype. Curr Opin Hematol 25, 212–218 (2018). 10.1097/MOH.0000000000000425

21 Kretschmer, M., Mamistvalov, R., Sprinzak, D., Vollmar, A. M. & Zahler, S. Matrix stiUness regulates Notch signaling activity in endothelial cells. Journal of Cell Science 136, jcs260442 (2023).

22 Rostama, B., Peterson, S. M., Vary, C. P. & Liaw, L. Notch signal integration in the vasculature during remodeling. Vascular pharmacology 63, 97–104 (2014).

23 Lobov, I. & Mikhailova, N. The role of Dll4/Notch signaling in normal and pathological ocular angiogenesis: Dll4 controls blood vessel sprouting and vessel remodeling in normal and pathological conditions. Journal of ophthalmology 2018, 3565292 (2018).

24 Liu, Z.-J. et al. Notch activation induces endothelial cell senescence and pro-inflammatory response: implication of Notch signaling in atherosclerosis. Atherosclerosis 225, 296–303 (2012).

25 Venkatesh, D. et al. RhoA-mediated signaling in Notch-induced senescence-like growth arrest and endothelial barrier dysfunction. Arteriosclerosis, thrombosis, and vascular biology 31, 876–882 (2011).

26 Miloudi, K. et al. NOTCH1 signaling induces pathological vascular permeability in diabetic retinopathy. Proceedings of the National Academy of Sciences 116, 4538–4547 (2019).

27 Cheng, Y.-L. et al. Evidence that neuronal Notch-1 promotes JNK/c-Jun activation and cell death following ischemic stress. Brain research 1586, 193–202 (2014).

28 Mo, J.-S. et al. Notch1 modulates oxidative stress induced cell death through suppression of apoptosis signal-regulating kinase 1. Proceedings of the National Academy of Sciences 110, 6865–6870 (2013).

29 Vilimas, T. et al. Targeting the NF-κB signaling pathway in Notch1-induced T-cell leukemia. Nature medicine 13, 70–77 (2007).

30 Kang, Y.-A., Pietras, E. M. & Passegué, E. Deregulated Notch and Wnt signaling activates early-stage myeloid regeneration pathways in leukemia. Journal of Experimental Medicine 217, e20190787 (2019).

31 Martínez-Zamudio, R. I. et al. AP-1 imprints a reversible transcriptional programme of senescent cells. Nature cell biology 22, 842–855 (2020).

32 Liu, Z. J. et al. Inhibition of endothelial cell proliferation by Notch1 signaling is mediated by repressing MAPK and PI3K/Akt pathways and requires MAML1. The FASEB journal 20, 1009–1011 (2006).

33 Netterfield, T. S., et al. Biphasic JNK-Erk signaling separates the induction and maintenance of cell senescence after DNA damage induced by topoisomerase II inhibition. Cell systems 14, 582–604. e510 (2023).

34 Coppé, J.-P. et al. Senescence-associated secretory phenotypes reveal cell-nonautonomous functions of oncogenic RAS and the p53 tumor suppressor. PLoS biology 6, e301 (2008).

35 Federman, N. Molecular pathogenesis of desmoid tumor and the role of γ-secretase inhibition. NPJ Precision Oncology 6, 62 (2022).

36 Pedrosa, A.-R. et al. Endothelial Jagged1 antagonizes Dll4 regulation of endothelial branching and promotes vascular maturation downstream of Dll4/Notch1. Arteriosclerosis, thrombosis, and vascular biology 35, 1134–1146 (2015).

37 Cherry, C. et al. Transfer learning in a biomaterial fibrosis model identifies in vivo senescence heterogeneity and contributions to vascularization and matrix production across species and diverse pathologies. GeroScience 45, 2559–2587 (2023).

38 Chung, L. et al. Interleukin 17 and senescent cells regulate the foreign body response to synthetic material implants in mice and humans. Science translational medicine 12, eaax3799 (2020).

39 Anderson, J. M., Rodriguez, A. & Chang, D. T. in Seminars in immunology. 86–100 (Elsevier).

40 Levenberg, S., Golub, J. S., Amit, M., Itskovitz-Eldor, J. & Langer, R. Endothelial cells derived from human embryonic stem cells. Proceedings of the national Academy of Sciences 99, 4391–4396 (2002).

41 Müller, A. M. et al. Expression of the endothelial markers PECAM-1, vWf, and CD34 in vivo and in vitro. Experimental and molecular pathology 72, 221–229 (2002).

42 Kim, I. L., Khetan, S., Baker, B. M., Chen, C. S. & Burdick, J. A. Fibrous hyaluronic acid hydrogels that direct MSC chondrogenesis through mechanical and adhesive cues. Biomaterials 34, 5571–5580 (2013).

43 Ahearne, M., Yang, Y., El Haj, A. J., Then, K. Y. & Liu, K.-K. Characterizing the viscoelastic properties of thin hydrogel-based constructs for tissue engineering applications. Journal of the Royal Society Interface 2, 455–463 (2005).

44 Visser, J. et al. Crosslinkable hydrogels derived from cartilage, meniscus, and tendon tissue. Tissue engineering part A 21, 1195–1206 (2015).

45 Hao, Y. et al. Dictionary learning for integrative, multimodal and scalable single-cell analysis. Nature biotechnology 42, 293–304 (2024).

46 Blighe, K., Rana, S. & Lewis, M. (Bioconductor, 2025).

47 Korotkevich, G., Sukhov, V. & Sergushichev, A. Fast gene set enrichment analysis. doi:10.1101/060012

